# Population structure determined by comparative amplicon gene sequencing in biofilm and planktonic bacterial communities from pesticide-contaminated water

**DOI:** 10.1101/705467

**Authors:** Jhenifer Yonara Lima, Cassiano Moreira, Gessica Costa, Paloma Nathane Nunes Freitas, Luiz Ricardo Olchanheski, Sonia Alvim Veiga Pileggi, Rafael Mazer Etto, Christopher Staley, Michael Jay Sadowski, Marcos Pileggi

## Abstract

Molecular and bioinformatics tools for research are very important for biodiversity characterization in contaminated environments due the presence of a high percentage of nonculturable microorganisms. The wide use of pesticides in agriculture exposes microbiomes to stressful and selective conditions that demand survival strategies such as biofilm formation. The purpose of this work was to evaluate the bacterial population structure in planktonic and biofilm communities of water that was used for washing the packaging of herbicides and stored for six months in tanks. This substance is highly contaminated waste. Samples of water and biofilms from tanks and biofilms developed for short times in flasks were used for DNA isolation and 16S rRNA gene sequencing. The physicochemical conditions imposed by water used for washing the containers were inadequate for killing the bacterial genera identified according to water and wastewater standards. The variation in population structure and diversity was lower in the planktonic samples than in the biofilm samples, indicating a possible combination of genetic drift and subsequent selection of individuals surviving under stressful water conditions, such as heating and contact with agrochemicals, over a six-month period. The biofilm formation in water tanks contaminated with pesticides enabled the survival of bacterial genera harboring the essential processes for adaptation to these environments; the presence of these processes was determined according to descriptions obtained from the genomic databases. This study suggests the potential of bacterial genera identified in biofilms obtained from tanks to adapt to contaminated environments through their metabolic complexity. Thus, herbicide biodegradation kinetics can be accessed through a culturable collection obtained from these communities.

## Introduction

Environmental microbiology has been gaining ground in academia over the last few years, bringing with it an interest in understanding composition, structure and function in nature [1]. Notably, contaminated environments expose microbiomes to stressful and selective conditions, and once microorganisms exhibit structural and metabolic adaptations, they can overcome these conditions and survive. These adaptive strategies are linked at the cellular level and to specific populations, allowing different responses to compounds such as pesticides [2,3] and enabling their use in biotechnological processes such as agrochemical degradation [4–6]

However, microbial diversity is high and poorly elucidated by culture techniques, even with different types of culture media, since a large percentage of these microorganisms are nonculturable [7,8]. Therefore, it is necessary to create new research tools that allow the characterization of microbiological biodiversity, i.e., molecular microbial ecology, through the use of high-throughput sequencing technologies with the aid of bioinformatics techniques for data analysis [1,9–11].

The knowledge and use of molecular technologies are fundamental to understanding the structure and functions of microbiomes and nonculturable microorganisms. Advances in genomics and metagenomics, marker genes, sequencing of the 16S rRNA gene and other molecular approaches are essential for predicting soil microbiome functions and improving agriculture production [12]. These approaches are fundamental to the creation of genomic libraries that allow us to identify and isolate enzymes with different biocatalytic activities from culturable strains, permitting their biotechnological exploitation [13,14]. These techniques allowed Krohn-Molt et al. (2017) to evaluate the genes of an association of algae essential to the carbon cycle with a unique and specific microbiome. Knowledge of the taxonomic structure of the microbiome increases the reliable handling of its latent biotechnological potential [16,17].

Thus, the composition and relative abundance of soil microbiomes in the Amazon were correlated with different types of land use, such as deforestation, which leads to carbon cycle disturbances and, consequently, climate change [18]. Therefore, microbiomes, which play key roles in biogeochemical cycles, are affected by anthropogenic action, particularly by the use of xenobiotic compounds, leading to changes in sustainability and environmental quality [18,19]. High-density DNA microarray and 16S rDNA amplicon next-generation sequencing were used to study the impact of the insecticide chlorpyrifos, the herbicide isoproturon and the fungicide tebuconazole on the soil bacterial community, and significant differences have been found in field experiments but not in soil microcosms exposed to these pesticides [20]. These methods were considered a guide for more specific studies in this area. A molecular approach was also used to analyze the effects of combinations of glyphosate and different herbicides on the rhizobacterial community of transgenic plants; this study indicated that glyphosate alone seems to be the less aggressive [21]. Molecular approaches also helped Dennis et al., 2018, reach the conclusion that the herbicides glyphosate, glufosinate, paraquat, and paraquat-diquat did not significantly interfere with soil microbial diversity and function under laboratory conditions.

In aqueous environments, which are also subject to contamination by agrochemicals, microbiomes present two possible lifestyles: planktonic and biofilm. The first term refers to the cells of microorganisms suspended in water (e.g., in an ocean water column) [22]. However, one of the main adaptive strategies found in microbiomes in contaminated or extreme environments is the formation of biofilms, in which cells adhere to a substrate or one another by secreting a gelatinous matrix of extracellular polymeric substances (EPS), which is configured as an extension of the cell itself [23,24]. The community can thus capture organic compounds for metabolization by extracellular enzymes through organized gene expression mediated by cell-cell communication or quorum sensing. This structure is also responsible for increased adaptability, stability, and cellular integrity [25–27]. 16S rRNA gene amplicon sequencing was used to verify the impact of herbicides and fertilizers on planktonic marine microbial communities at the Great Barrier Reef. Water characteristics, such as salinity, rainfall, temperature and water quality, have a great influence on the composition of microbial communities [28].Aquatic environments are complex and subject to various physicochemical interferences, making it difficult to understand the impact of agrochemicals on microbiomes.

Thus, the objective of this study was to identify the population structure of planktonic and biofilm bacterial communities in an environment consisting of tanks used for storing washing water from the packaging of herbicides, which results in highly contaminated waste; thus, another objective was to glimpse adaptive strategies through the composition of genes present in their taxonomic units.

## Materials and methods

### Study area

The object of study was water used to wash the containers (Fig 1) of 30 agrochemicals (S1 Table). The water used for the washing of agrochemical packages came from an artesian well. After washing, the water was kept in two 10,000 L tanks for 6 months at the Capão da Onça Farm School, State University of Ponta Grossa, State of Paraná, Brazil (25°05’31.3“S; 50°03’28.0“W) (altitude: 1000 m). The first tank (T1) had water from the washing of agrochemical containers that had lost some volume by an evaporation process at 100 °C for 190 min and was then stored in a second tank (T2) until being transported and discarded.

**Figure 1:**
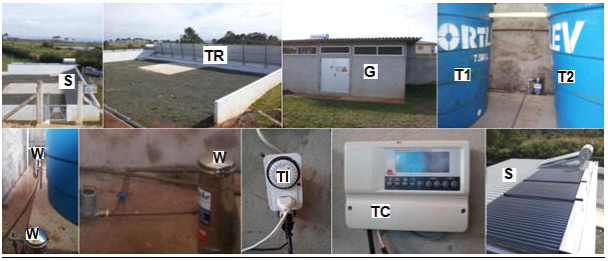
Evaporation of the water from the washing of pesticide containers used for DNA collection in this study. S: Sanitary sink; TR: Tractor spray washing trough; SH: Shed where T1 and T2 are; T1: Tank 1; T2: Tank 2; W: Water filters; TI: Timer for controlling the flow of water; TC: Solar heating temperature control panel; So: Solar heating system.

### Physicochemical analyses of water

The following physicochemical characteristics of water samples collected from the tanks and the artesian well were analyzed according to the *Standard methods for the examination of water and wastewater* (1998): chlorine, conductivity, color, fluorine, pH, total dissolved solids and turbidity, biochemical oxygen demand (BOD) and chemical oxygen demand (COD).

### Collection of samples and extraction of DNA

Four liters of water were collected from T1 and T2 to obtain planktonic samples. The water was passed through autoclaved glass fiber membranes in 750 mL triplicates: first, a 5-μm-thick prefilter; then, 1.2 μm; and last, 0.45 μm (Merck Millipore, Carrigtohill, Ireland). After the last membrane, the samples were stored at −80 °C. The samples were named M1 (preheating) and M2 (postheating).

The biofilm samples were collected in triplicate from the wall surface of the tanks in contact with the water line using 3 sterile swabs for each triplicate. The samples were named B1 (preheating) and B2 (postheating). Likewise, biofilms were collected from the 4 L flasks in which water samples were collected after being kept immobile for five days. The samples were named G1 (preheating) and G2 (postheating). The material collected in triplicate was stored at −80 °C.

Samples of planktonic origin (retained in the membranes) were macerated in liquid nitrogen with the aid of a pestle. The swabs used for the biofilm collections were submerged in 1 mL of saline buffer (0.9% NaCl) and homogenized for 5 min to extract biological material from the cells in the buffer. DNA extractions were performed using the PowerSoil DNA extraction kit (MO BIO Laboratories Inc., Carlsbad, USA), and the material obtained was stored at −80 °C.

### Sequencing

Sequencing of the samples was performed using 515F/806R primers, which are specific for the V4 region [29]. Amplification and sequencing were performed using the dual index method reported by the University of Minnesota Genomics Center (Minneapolis, MN, USA) [30]. A negative control containing sterile water was included in each amplified and sequenced sample plate. Sequencing was performed on the Illumina MiSeq platform using an iSeq 100 System (Illumina, Inc. San Diego, USA) [29]. Sequence data are available from the Sequence Read Archive of the National Center for Biotechnology Information under accession number SRA: PRJNA528924.

### Bioinformatics

Sequence processing was performed using mothur software (version 1.29.2). The prokaryotic sequence data were trimmed to the first 160 nucleotides (nt), and end pairing was carried out using fastq-join software [31]. The sequences were trimmed for quality following a previous description for V5-V6 data [32]. The overall alignment was performed in relation to the SILVA database ver. 119 [33], the sequences were subjected to a 2% preclustering to remove sequence errors [34], and the chimeric sequences were identified and removed by UCHIME software [35]. Operational taxonomic units (OTUs) were assigned at ≥ 97% identity by complete linkage clustering. Taxonomic attributions were made by comparing the Ribosomal Database Project ver. 14 to a bootstrap limit of 60%, as previously described [36].

Bacterial communities were ordered based on OTU abundance using multidimensional scaling. The relative abundance of the taxonomic groups and the functional abundances inferred in PICRUSt were regressed in relation to the ordering scores to determine significant correlations with the ordering scores of the bacterial communities [37]. The 16S rDNA data were analyzed as indicated by the PICRUSt genome prediction software [http://picrust.github.io/picrust/] from raw sequence reads in the following environment: NumPy (1.7.1), biom-format (1.3.1), PyCogent (1.5.3), PICRUSt (1.0.0-dev), and PICRUSt script (1.0.0-dev). Functional predictions were assigned up to KO tier 3 for all genes. Results from the MG-RAST QIIME report were compared with predictions from PICRUSt using the “compare_biom.py” subroutine with normalization and observations not in the “expected data” file ignored. Results were compared using both shotgun metagenomic and PICRUSt predictions as the expected data. Statistical analyses were performed using the vegan package in R [38].

## Results and discussion

This work discusses three levels of results to account for interferences in the adaptive strategies presented by the different bacterial communities linked to the contaminated environment in this study: the physicochemical characteristics of the environment, the taxonomic composition at the time of collection, and the genes associated with the identified taxonomic units.

### Physicochemical characteristics of the tank water

The physicochemical data are analyzed in this section because they provide the conditions under which the adaptation of the present microbiome to the environment occurs. The data for the water from T1 and T2 and the artesian well whose water was used in the washing are shown in Table 1.

**Table 1:** Physicochemical analyses of tanks 1 (T1) and 2 (T2) and the artesian well (AW). Parameter values are presented as the means of triplicate measurements. WHO: World Health Organization; SD: standard deviation; TDS: total dissolved solids; BOD: biochemical oxygen demand; COD: chemical oxygen demand; ND: not detected. The letters A, B, C, and D represent significant differences within each row.

| Parameters | T1 | SD | T2 | SD | AW | SD | WHO |
| --- | --- | --- | --- | --- | --- | --- | --- |
| Chlorine (mg/L) | 1.66<sup>A</sup> | 0.1 | 1.57<sup>A</sup> | 0 | 0<sup>B</sup> | 0 | 0.7<sup>C</sup> |
| Conductivity (μS/cm) | 418.3<sup>A</sup> | 2 | 413<sup>A</sup> | 0.7 | 13.57<sup>B</sup> | 0.564 | 10 – 100<sup>B</sup> |
| Color (Pt/Co) | 245<sup>A</sup> | 1.5 | 250<sup>B</sup> | 0 | 0<sup>C</sup> | 0 | 0<sup>C</sup> |
| Fluorine (mg/L) | 0.82<sup>A</sup> | 0 | 0.81<sup>A</sup> | 0 | 0.05<sup>B</sup> | 0 | 1.5<sup>C</sup> |
| pH at 25°C | 5.45<sup>A</sup> | 0.1 | 5.32<sup>A</sup> | 0 | 4.75<sup>B</sup> | 0.045 | 6.5 – 8.5<sup>C</sup> |
| TDS (ppm/NaCl) | 186.3<sup>A</sup> | 1 | 183.2<sup>A</sup> | 0.3 | 0<sup>B</sup> | 0 | 100 - 600<sup>A</sup> |
| Turbidity (NTU) | 204<sup>A</sup> | 5.8 | 206.3<sup>A</sup> | 3.6 | 0.25<sup>B</sup> | 0 | 1<sup>C</sup> |
| BOD | 80.3<sup>A</sup> | 1.5 | 69<sup>B</sup> | 4.6 | ND | ND | 0<sup>C</sup> |
| COD | 1352.3<sup>A</sup> | 29 | 1105<sup>B</sup> | 35.5 | ND | ND | 0<sup>C</sup> |

According to the parameters shown in Table 1, the water characteristics of tanks 1 and 2 are not significantly different from each other.

The data obtained in this work indicate that the heating process did not interfere with the physicochemical parameters of the water in the tanks. Nevertheless, there are significant differences between tank water and the water from the artesian well in every analyzed item, mainly due to the presence of pesticides that significantly altered the characteristics (Table 1) and independent of the microbiome of these communities.

The chlorine and fluorine contents of T1 and T2 are noteworthy since, according to the World Health Organization [39], these specific values are close to the acceptable concentrations for drinking water, even though the water from the artesian well and that from the abovementioned tanks had not been added to these samples (Table 1). Moreover, according to the WHO, the pH is also at inadequate levels for bacterial growth, since the pH values are close to that of drinking water. In addition to these parameters, the increase in conductivity and COD indicated important changes in the physicochemical characteristics of the water of both tanks from those of water from the artesian well, most likely related to the nonbiological oxidation processes of the pesticides. The BOD data indicate biological activity, even in an environment unfavorable to the development of microorganisms. COD values are usually higher than BOD values due to the greater ease of compound oxidation by the chemical route than by microorganisms [40].

The physicochemical data obtained in our study were compared to those of water with different origins and from environments with different characteristics to determine which water the T1 and T2 waters most resemble. Such environments were that of drinking water [39], the interior transition zone of the ocean [41], the Red Sea [42], southwest part of the Gulf of Mexico [43], a hot water spring in Siloam, South Africa [44], an integrated system of anaerobic-aerobic reactors for the treatment of wastewater (WW) in Ethiopia [45] and the Yeongsan River Basin of South Korea [46]. The two tanks do not significantly differ in chlorine level, but these levels are statistically higher than those found for seawater and potable water (Table 2). However, the levels found for the well water are statistically similar to those of the sea. Fluorine levels are statistically similar in both tanks; however, they are lower than those of drinking water and higher than those of the artesian well water (Table 2) [44]. The tanks have statistically similar pH values but are more acidic than the artesian well, residual tributaries, ocean, fresh water and drinking water and are statistically significantly different from these sources in pH (Table 2). In general, when total dissolved solids (ppm/NaCl, corresponding to salinity) are analyzed, our data present statistical similarities to the levels in WW, drinking water and a hot water spring in Siloé, South Africa. The BOD and COD rates are higher in T1 than in T2, indicating that there are probably more contaminants and microorganisms in the former. For the same reason, the rates in T1 and T2 are higher than those in the artesian well. The tanks have lower BOD and higher COD than WW and a wastewater tributary (WT), probably because the methods used in the treatment of these effluents have microorganisms that increase their BOD values but not their COD values in relation to their values in the tanks studied in this work (Table 2).

**Table 2:**
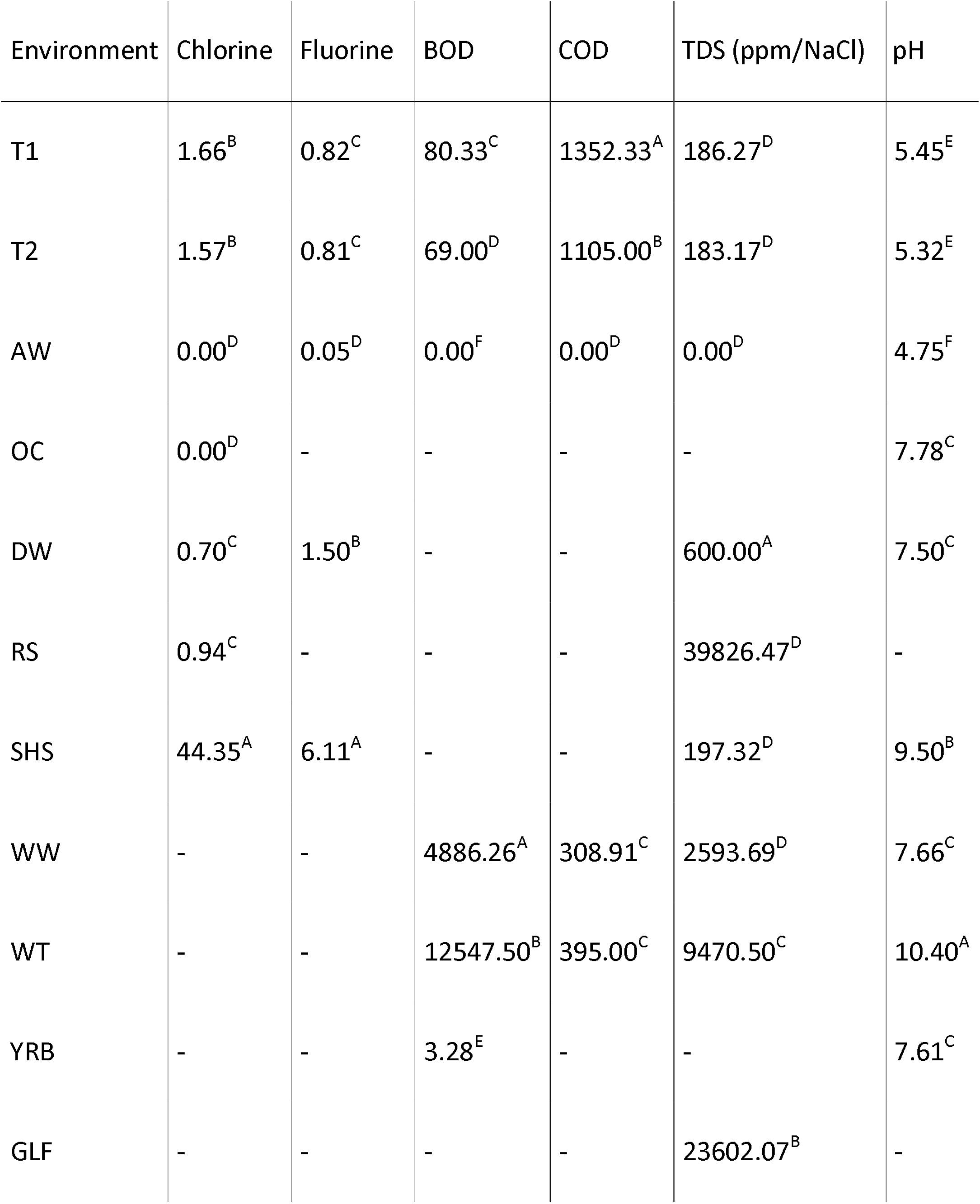
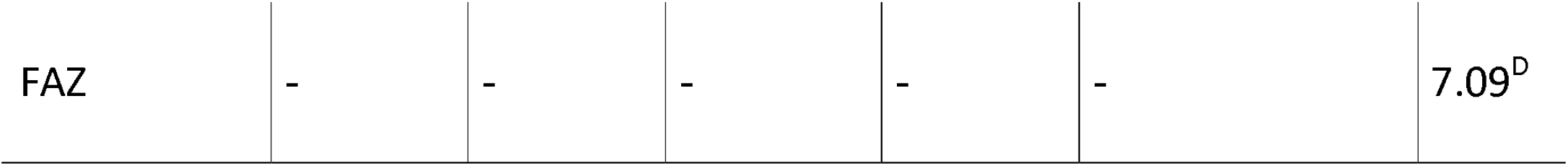
Comparison of physicochemical analyses of tanks 1 (T1) and 2 (T2) and the artesian well (AW) with different environments. Data were obtained in the literature for environments such as the oceanic transition zone (OC), drinking water (DW), the Red Sea (RS), Siloé Hot Spring (SHS), wastewater (WW), wastewater tributary (WT), the Yeongsan River Basin (YRB), the Gulf of Mexico (GLF) and freshwater aquaculture systems (FAS). No information (−). The letters A, B, C, D, E, and F represent significant differences within each column.

In this way, tanks T1 and T2 were characterized as environments with inadequate levels of chlorine and fluorine for the survival of the genera identified in this study. However, these genera were probably able to tolerate these conditions, either by inactivity in planktonic form or by organic and metabolic activity in biofilms [47].

It is important to remember that these physicochemical analyses reflect the planktonic community more directly than they do biofilms. Biofilms, although interfacing with the environment, most likely present modifications that improve metabolism and the adaptability of microorganisms to the studied environment [26], as will be discussed next.

### Biodiversity and bacterial taxonomy

The identification of the taxonomic groups present in the different niches evaluated in this study may indicate changes in bacterial population structures adapting to the stress produced by the physicochemical conditions and contamination. Sample coverage data in terms of the Simpson indices from the 16S rRNA gene sequencing and diversity calculations are shown in Fig 2.

**Figure 2:**
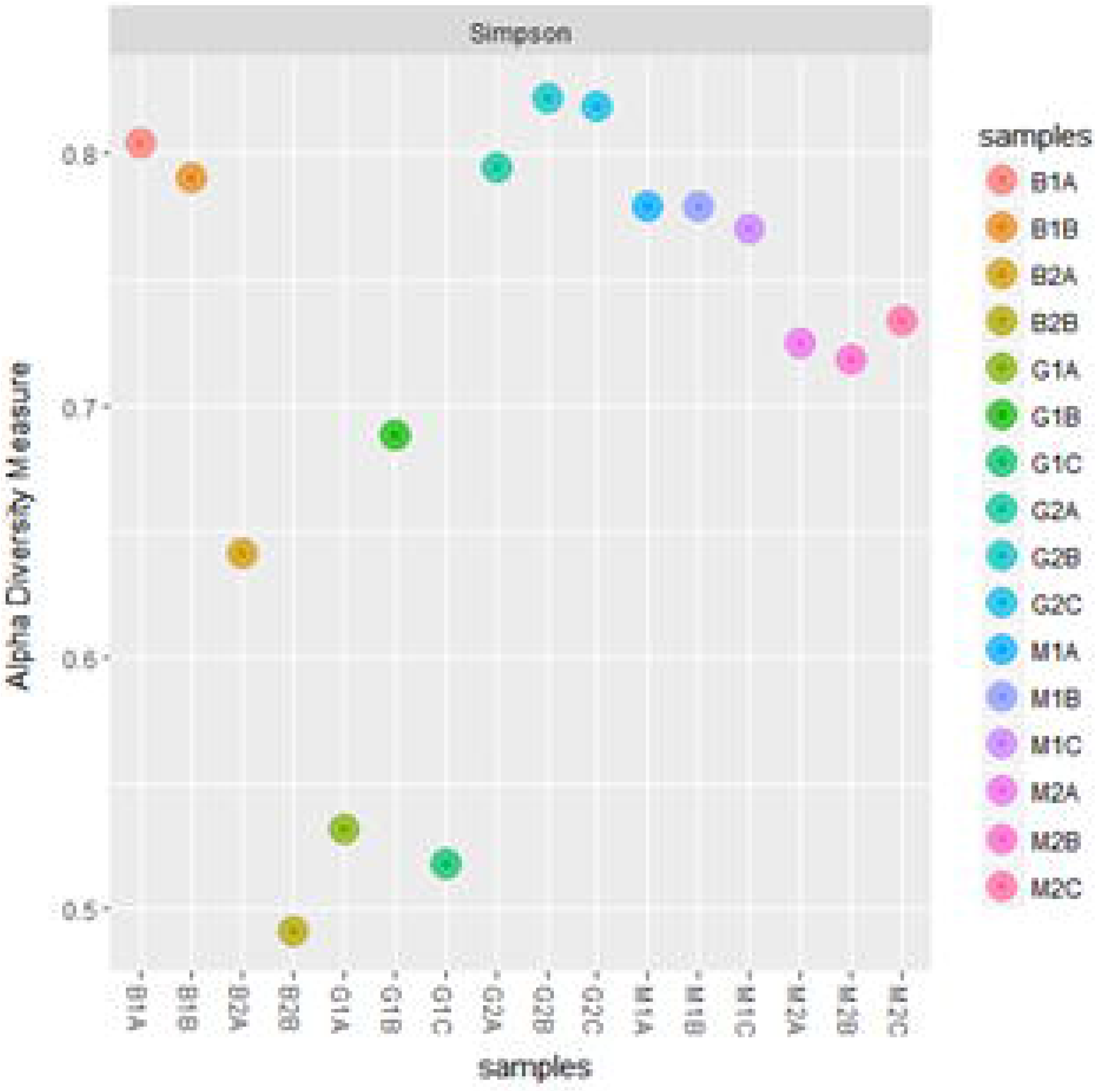
Measurements of alpha diversity in the planktonic triplicate (M1A, M1B, M1C and M2A, M2B, M2C) and biofilm triplicate (B1A, B1B and B2A, B2B) samples from the tanks and biofilm samples from the container’s triplicates (G1A, G2B, G2C and G2A, G2B, G2C).

Analyses of the sequences obtained in every community studied in this work showed that the most representative classes are Gammaproteobacteria, Alphaproteobacteria and Betaproteobacteria, as shown in Fig 3.

**Figure 3:**
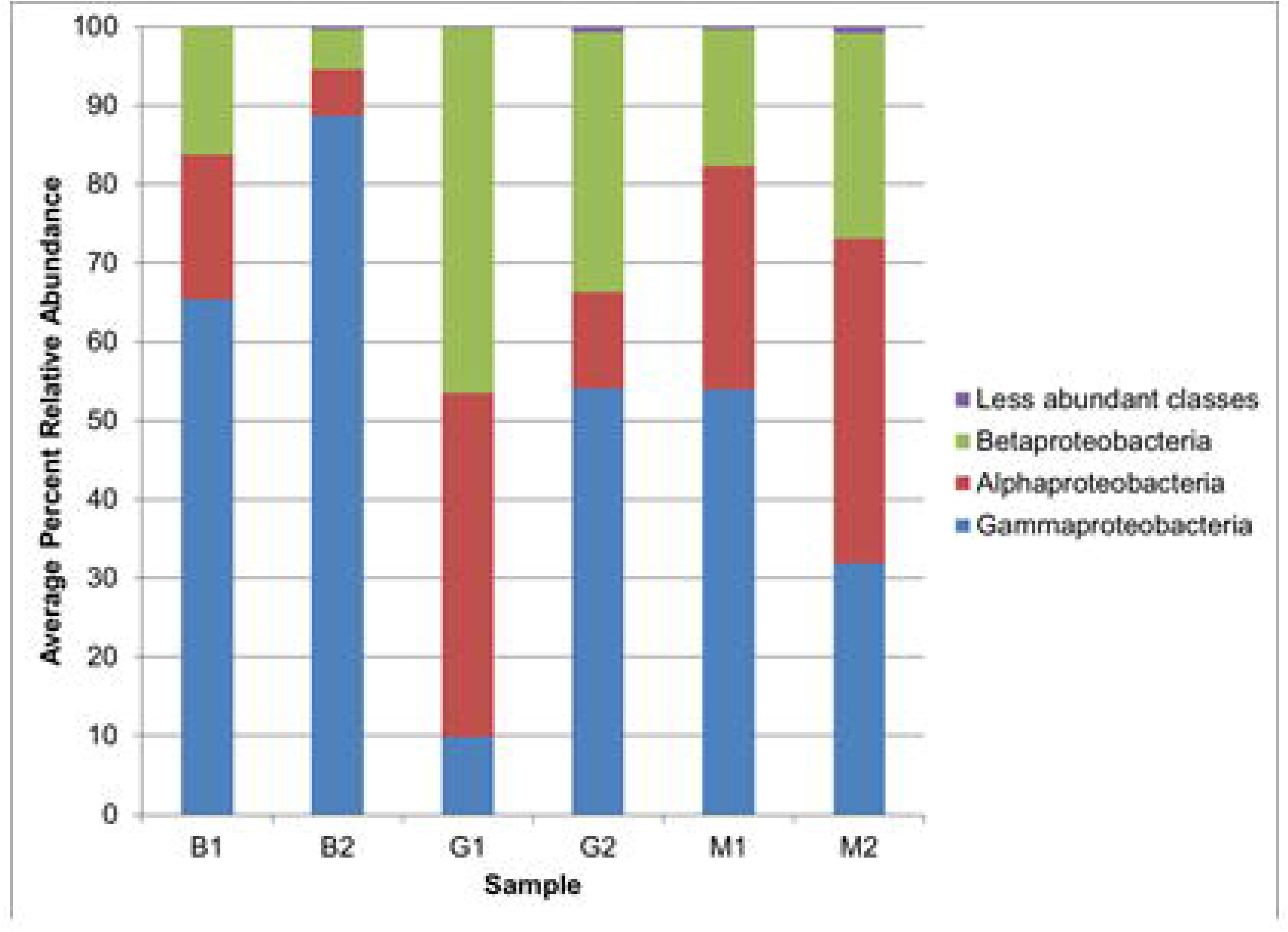
Abundance of bacterial classes in the biofilm samples from the containers and planktonic samples from tanks 1 and 2. B1 and B2: taxonomic units identified from pre- and postheating biofilms, respectively; M1 and M2: taxonomic units identified from pre- and postheating planktonic samples, respectively; G1 and G2: taxonomic units identified from biofilms formed in containers with pre- and postheating water, respectively.

The most representative families identified by the sequences obtained in every community are shown in Fig 4.

**Figure 4:**
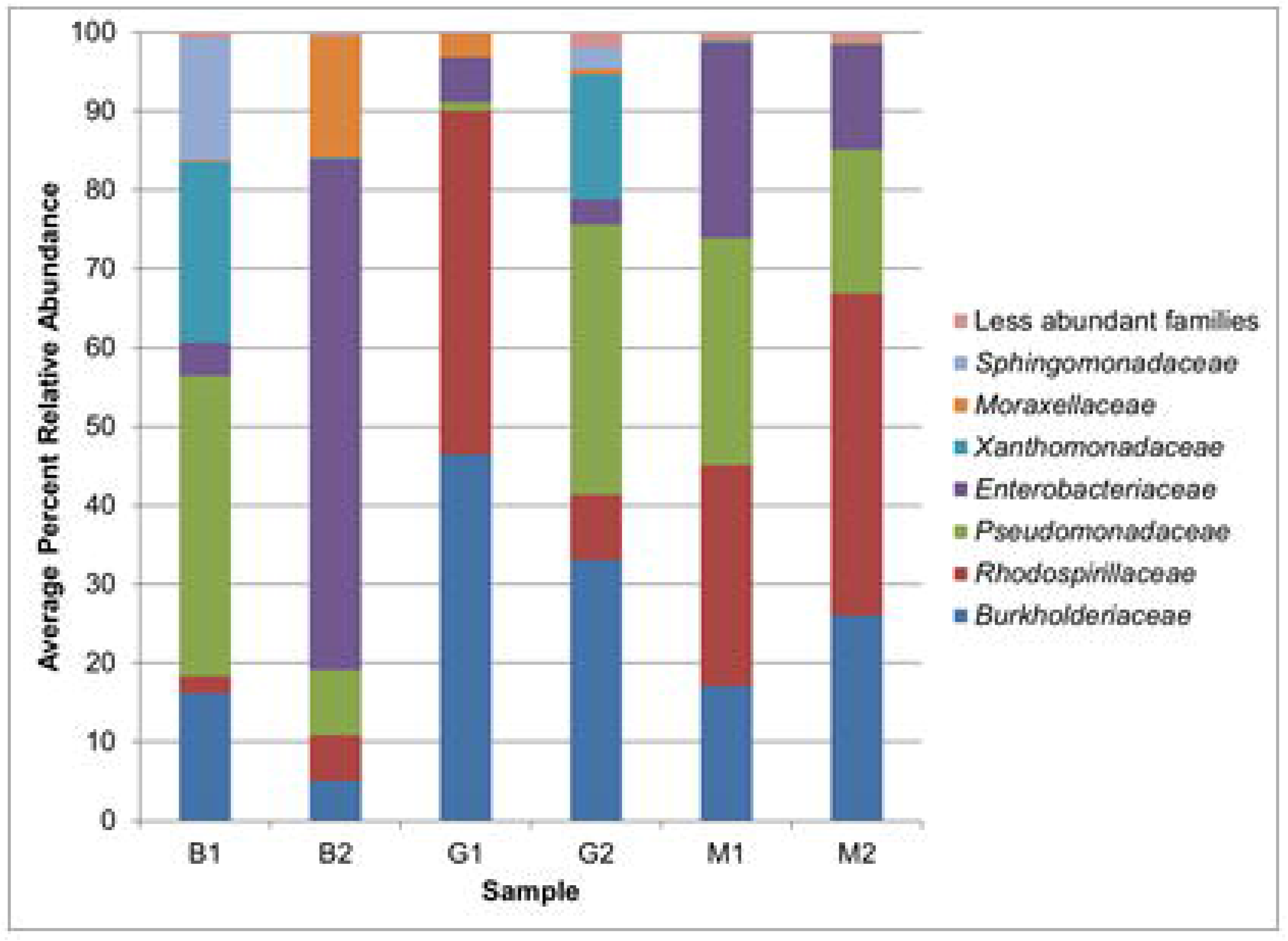
Abundance of bacterial families in biofilm samples from tanks 1 and 2. B1 and B2: taxonomic units identified from pre- and postheating biofilms, respectively; M1 and M2: taxonomic units identified from pre- and postheating planktonic samples, respectively; G1 and G2: taxonomic units identified from biofilms formed in containers with pre- and postheating water, respectively.

The number of sequences assigned by community and by taxonomic category and their respective percentages within each community are shown in S1 Fig.

Simpson’s diversity index demonstrates the distribution of alpha diversity (Fig 1). The sequences obtained and identified as the genera of the planktonic communities (M1 and M2) are represented in the biofilm communities (B1 and B2, G1 and G2) but have different diversity indices that depend on selection and adaptation to the environment (Figs 1 and S1 Fig).

The data shown in Figs 3 and 4 indicate that there is great variation in the population structure in different communities in relation to classes and families. The event of water heating is a nondirectional selective process since none of the taxonomic groups identified in this study have resistance responses for high temperatures, but it changes the number of taxonomic units, resulting in decreased diversity in the postheating samples for every community and less intense changes for the planktonic communities than for the biofilm communities. The effect of this genetic drift that began in the planktonic community can be observed in Fig 2 and leads to different diversity and abundance among the communities that settled in the biofilm structure in the tanks and in the flasks before and after the heating process. The planktonic community has a direct influence on biofilm formation [48]. Chronologically, the communities are formed in the following order: first T1, followed by B1 and then G1. This ordering is valid for postheating communities, with the proviso that the T2 community is formed after the T1 community.

A genetic drift effect is more conspicuous in the biofilm population structures in flasks G1 and G2, formed from the planktonic strains, than in the structures in other communities. This is probably because they had less time to consolidate population structures that are more adapted to the environment, as observed in B1 and B2.

Population structure variation is lower in the planktonic samples than in the biofilm samples in tanks and flasks, which may indicate that there was a combination of the genetic drift of individuals who survived the heating and subsequent selection of differentiated characteristics to allow individuals to adapt to the tank environment. This environment for water contaminated with 30 different agrochemicals and with physicochemical conditions unfavorable to the survival of microorganisms identified in this work was considered stressful (Table 1 and S1 Table). The organization of bacteria in biofilms in tanks and containers, with their cooperative or antagonistic interactions, may lead to a specific combination of species with increased adaptability to the environment [49]. Thus, the most dominant sequences in B2, corresponding to the genera *Enterobacter* and *Acinetobacter*, are poorly represented in the other communities studied, including the biofilms in B1, G1 and G2 (Fig 2).

Theoretically, the transition from planktonic to biofilm life involves changes in the regulation of different groups of genes and can be seen as a transition phase before the colonization of a new environment [50]. Therefore, this transition should not present significant changes in the population structure (Fig 3). Throughout the establishment of the community, the most abundant genera tend to retain most of the genes related to basic cellular processes, leading to the same function being shared by individuals of different species and not only being distributed by the selection of the fittest [51].

Nevertheless, the reduction in diversity after the water heating event possibly led to genetic drift, altering the fine structure of the M2, G2 and B2 in relation to that of the M1, G1 and B1 communities and the retention of genes involved in central adaptive processes. If the genetic drift and the functional diversity of different bacterial communities were less preponderant, the diversity indices of B1 and B2, B2 and G2, and G1 and G2 should be closer [48]than what was observed in Fig 2. In addition, the data shown in S1 Fig indicate that the same genus but with different 16S sequences may contribute to different communities adapted to the environments studied in this work [52]. Thus, the genetic drift hypothesis was established from the sequences identified and not by the taxonomic units.

Aquatic environments are complex [28]; therefore, the analysis of changes in the structure of communities of microorganisms that are made to adapt to environmental contaminants is difficult. Restricted environments with a high number of agrochemicals, such as those presented in this paper, allow basic hypotheses to be tested with increased safety before being tested at a larger environmental scale.

### Functional diversity

The comparative data in Fig 5 cluster genes by the sequences of the 16S rRNA gene identified in this study. This analysis was carried out as a survey of biotechnological potential [18,19] mainly related to biodegradation in the storage tanks of water contaminated with agrochemicals, which are considered a significant environmental liability.

**Figure 5:**
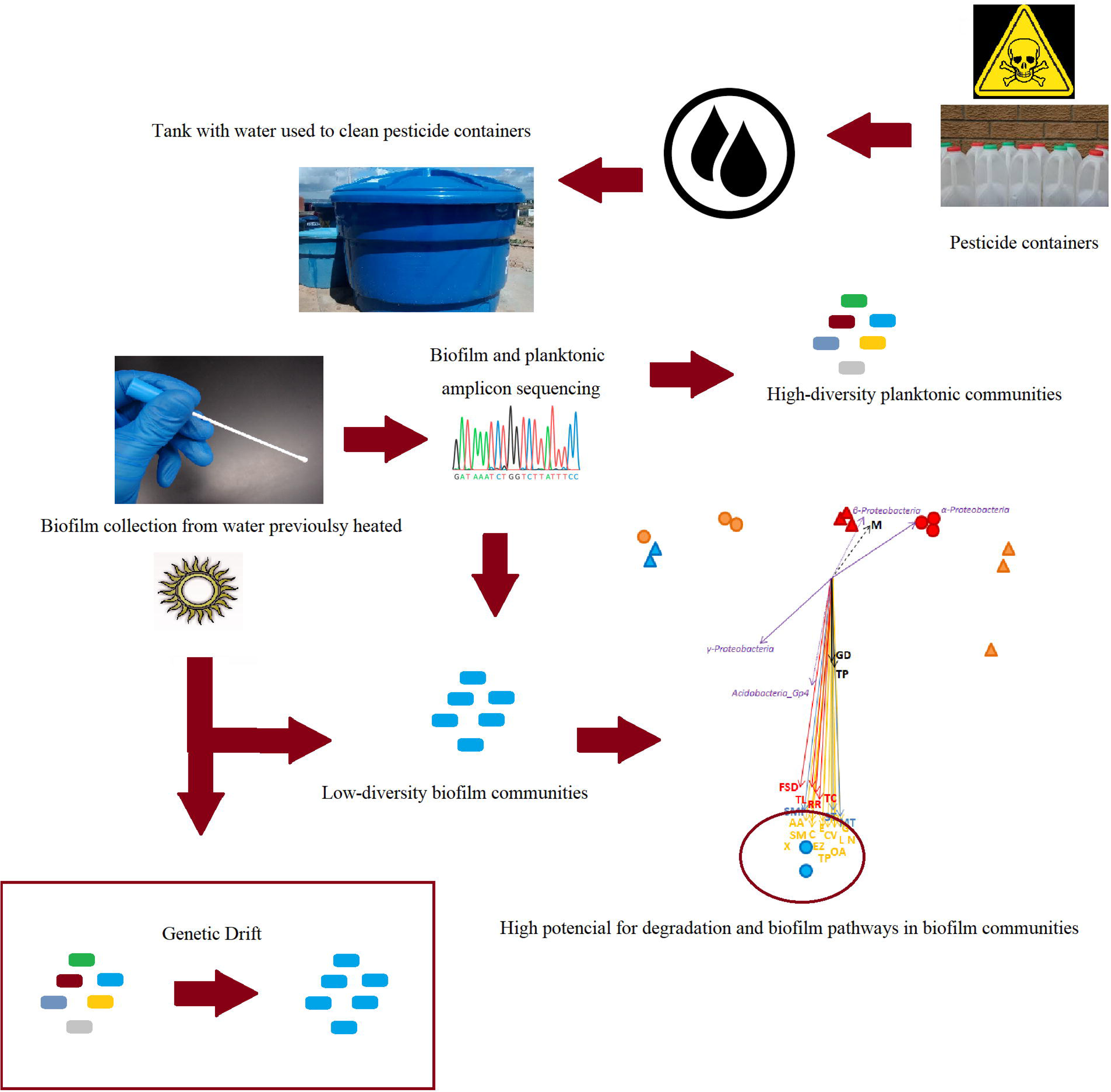
Ordering the sites based on OTU abundance by multidimensional scaling. Only the bacterial genus (green), genetic process (red), cellular process (black), environmental process (blue) and metabolic pathways (yellow) with p<0.05 after 999 permutations are displayed. Each vector shows the direction of increase for a given variable, and its length indicates the strength of the correlation between the variable and the ordination scores. Legend: M: Motility; GD: Growth and Death; TP: Transportation in Catabolism; MT: Membrane Transport; SMI: Signaling Molecule Interaction; ST: Signal Transduction; TL: Translation; TC: Transcription; RR: Replication and Repair; FSD: Folding, Sorting and Degradation; AA: Amino Acid; SM: Secondary Metabolites; C: Carbohydrate; E: Energy; EZ: Enzyme; G: Glycan; L: Lipid; CV: Cofactors and Vitamins; OA: Other Amino Acids; TP: Terpenoids and Polyketides; N: Nucleotide; XB: Xenobiotic Biodegradation.

Nonmetric multidimensional scaling (NMDS) shows the correlations between the number and types of 16S rRNA gene sequences grouped by the 6 different communities. Genes related to bacterial motility have a high correlation with the two planktonic communities (S2 Table).

On the other hand, genes associated with other cellular processes, environmental processes, gene information processes and metabolic processes have a strong correlation with the biofilm community B2 (Table 1), which is predominantly composed of the genera *Acinetobacter* and *Enterobacter*.

To picture the potential for the degradation and bioremediation of pesticides and relate the hypotheses about adaptation to the environment described in this study, the degradation capacity of the pesticides (S1 Table) was related to the genera of the most prevalent bacteria identified by the 16S rRNA gene sequencing (Fig 2).

Bacteria from the genus *Pseudomonas* (representing 45% of the structure in B1 and 8% of the structure in B2, Fig 4) present in communities identified in Hawaiian soil can degrade pyraclostrobin, which is present in several fungicides [53]. A strain of *Pseudomonas aeruginosa* isolated from rice-cultivated soil has the capacity to degrade propiconazole [54]. Other strains of the genus *Pseudomonas* isolated from agricultural soil degrade thiamethoxam [55].

Strains of the genus *Enterobacter* that were identified in Egyptian soil (representing 63% of the structure in B2, Fig 4) degraded the herbicide Topik [56]. *Pseudomonas* and *Acinetobacter* (representing 15% of the structure in B2, Fig 4) from communities where there was tar processing were identified as potential degraders of mineral oil [57]. A strain of the genus *Burkholderia* was isolated from paddy fields with pesticide application in Koppal, India, and was a degrader of propiconazole, a component of the fungicide Tilt [54].

The genera in the literature that are most related to agrochemical biodegradation are the same as those that dominate B2. Thus, the genera identified in this study have agrochemical degradation potential and the capacity for structuring biofilm communities, as shown in Fig 2, suggesting the viability of searching for promising strains among the culturable collection obtained in this study. The high temperature to which the planktonic community was subjected is bactericidal for any genus observed in this study [39]. Therefore, the nondirectional selective effect induced by the high temperature was independent of the genomes of the identified genera.

### Comparative 16S rRNA gene sequencing

In this work, a model was proposed to explain a possible range of characteristics whose expression could confer adaptive advantages over the stressful environment of water tanks used in the washing of agrochemicals. The population fluctuations observed were the result of genetic drift and the capacity for biofilm formation, which involves membrane transport, interaction of signaling molecules and signal transduction, and functional bacterial versatility [58]. The biofilm structure depends on the presence of EPS, which involves the production of glycans and lipid transportation in catabolism. Biofilms confer genomic plasticity and phenotypic flexibility to the species that compose them, providing successful reproduction in stressful environments (Fig 5)[23].

Theoretically, these two characteristics, plasticity and flexibility, and community diversity, which are higher in the planktonic community (Fig 2) than in the biofilm community, may lead to species fixation over time or even extinction of the less efficient. Some mutations may be randomly lost by genetic drift; however, significant advantageous mutations are more likely to fix in the population, as is the case of motility in a planktonic community, the only feature that has a high correlation with this community (Fig 5). If the selection pression in a community is strong, the frequencies of genotypes that improve competence increase in the population [59]. This model can be applied for mutations in populations with one species or populations of different species but with gene flow potential, as might occur in the B2 community.

The environments studied in this work passed through a sequence of physicochemical changes involving chemical contamination and heat over six months. During these events, genera that had the capacity to develop useful functions [58] to consolidate biofilms, such as *Enterobacter* spp. and *Acinetobacter* spp., dominated the B2 community [48,60], which was formed after the communities M1, M2 and B1 by selection and genetic drift.

Environmental biofilms have a level of metabolic complexity higher than that of their individual components [61]. Thus, the study of biofilm formation in environments contaminated with pesticides, which can be stressful to microorganisms, has the potential to produce information about response systems for adaptation. This information provides a possible way to exploit enzymatic potentials, mainly those related to biodegradation and bioremediation of herbicides [50], in the kind of waste used in this study. New knowledge can be obtained for the exploitation of the enzymatic potential of bacterial communities under these environmental conditions, mainly the potential related to the biodegradation kinetics of herbicides.

## Conclusions

The population structure and abovementioned functional profile derived from amplicon sequencing of bacterial communities from tanks that contained water used for washing the packaging of herbicides and that had been stored for six months to form a highly contaminated waste were studied in this work. The communities were characterized as tank biofilms, flask biofilms, and planktonic samples, all from pre- and postheating treatments and formed at different periods of time. Only two genera identified in this study, *Acinetobacter* and *Enterobacter*, were dominant in the tank biofilms after water heating, which probably acted as a genetic drift agent for the planktonic community. A database was accessed after the amplicon sequencing to compare genomic compositions, and this analysis showed that these genera have a metabolic potential for survival under these environmental conditions by possessing genes such as those related to biofilm formation, structure and membrane transport, quorum sensing and xenobiotic degradation. The genetic drift, induced by water heating, decreased the diversity of the bacterial biofilm community, which probably survived as large bacterial populations under the harmful conditions found in the tank due their genomic diversity and adaptation to environmental responses, since it is known that biofilms have a level of metabolic complexity higher than that of their individual components. Thus, the hypotheses about metabolic abilities, generated for a small and controlled environment as that studied in this work, can be tested with communities isolated from the biofilms formed in these tanks. Thus, new knowledge can be obtained for the exploitation of the enzymatic potential of bacterial communities, mainly the potential related to the biodegradation kinetics of herbicides.

## Supporting information

Cover Letter Plos One

English Review Certificate

## Acknowledgments

The authors thank Maria Janina Pinheiro Diniz for assisting in the preparation and execution of experiments.

## Supporting information

**S1 Fig:**
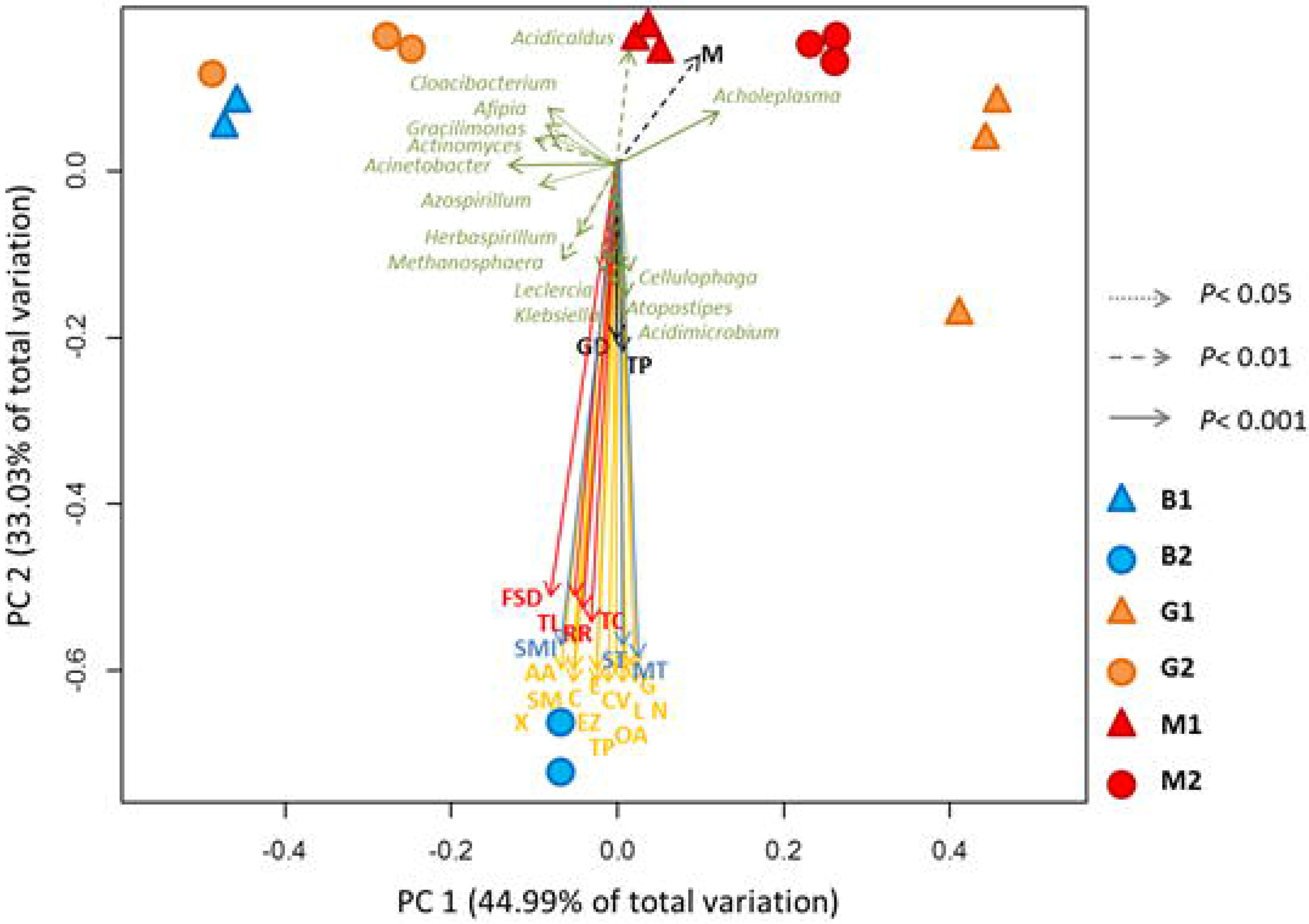
Number of sequences assigned by community and by taxonomic category and their respective percentages within each community. M1: preheating planktonic community; M2: postheating planktonic community; B1: preheating biofilm community; B2: postheating biofilm community; G1: preheating container biofilm community preheating; G2: postheating container biofilm community.

**S1 Table:** Agrochemicals whose containers were washed and the water used in the process to obtain the samples evaluated in this study.

**S2 Table:** Correlations of the functional characteristics identified in our analyses of the studied communities. M1: preheating planktonic community; M2: postheating planktonic community; B2: postheating biofilm community; r2: coefficient of determination; p: coefficient of significance.

