## Supplementary figures and images for "Population structure determined by comparative amplicon gene sequencing in biofilm and planktonic bacterial communities from pesticide-contaminated water"

### Supplemental Figure 1

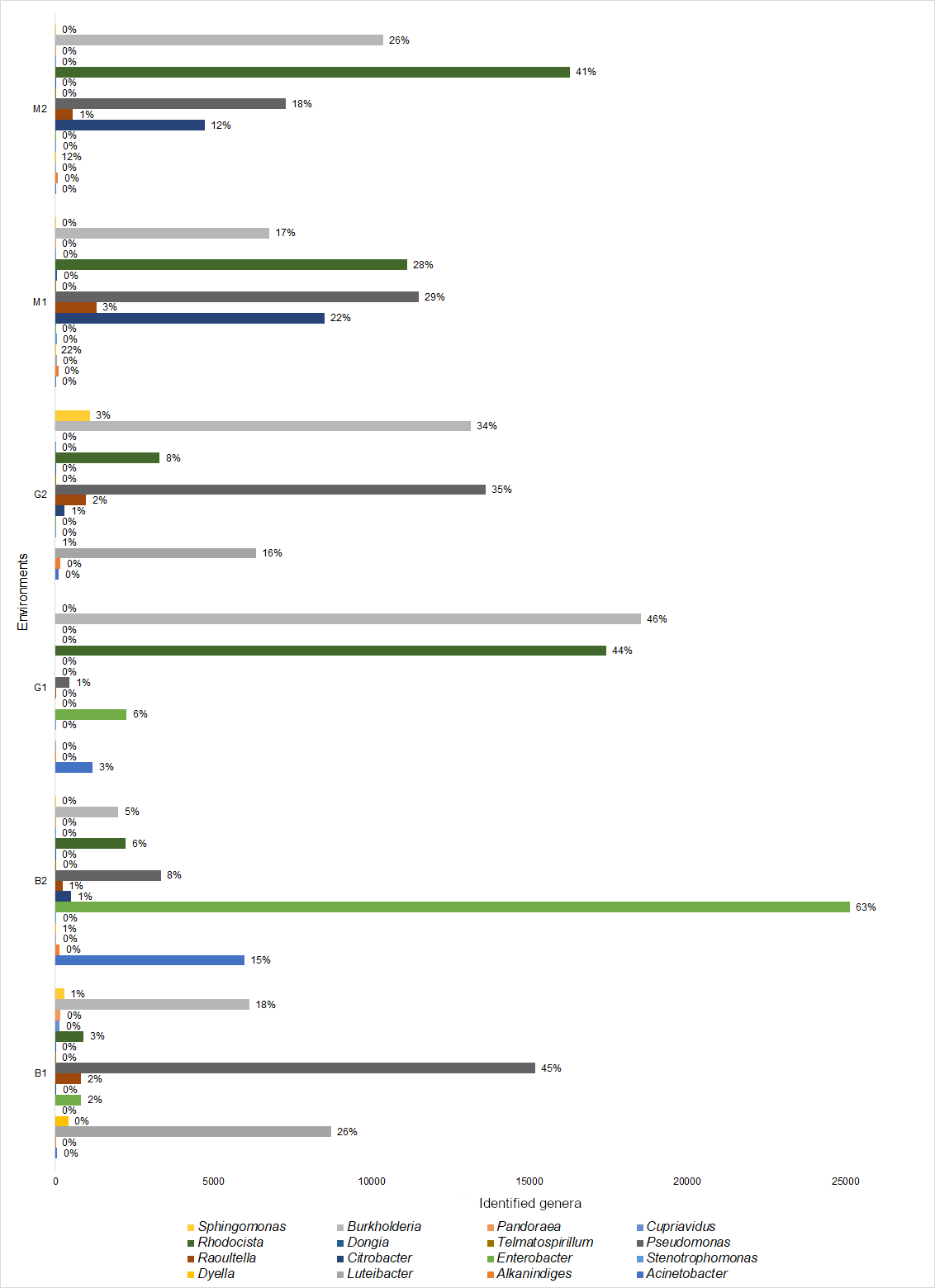
