## Supplemental Table 1 for "Population structure determined by comparative amplicon gene sequencing in biofilm and planktonic bacterial communities from pesticide-contaminated water"

| Product | Function | Manufacturer | Composition |
|---|---|---|---|
| **FUNGICIDES** | | | |
| Abacus HC | Fungicide | BASF | Pyraclostrobin, epoxiconazole |
| Ativum EC | Fungicide | BASF | Epoxiconazole, fluxapyroxad, pyraclostrobin |
| Brio | Fungicide | BASF | Epoxiconazole, kresoxim-methyl |
| Caramba 90 | Fungicide | BASF | Metconazole |
| Comet | Fungicide | BASF | Pyraclostrobin, epoxiconazole |
| Corbel | Fungicide | BASF | Fempropimorph |
| Difere | Fungicide, bactericide | Oxiquímica Agrociência LTDA | Copper oxychloride, metallic copper equivalent |
| Forum Plus | Fungicide | BASF | Dimethomorph, chlorothalonil |
| Fox | Fungicide | Bayer S.A. | Trifloxystrobin, prothioconazole |
| Opera SE | Fungicide | BASF S.A. | Pyraclostrobin, epoxiconazole |
| Opera Ultra | Fungicide | BASF S.A. | Pyraclostrobin, metconazole |
| Orkestra SC | Fungicide | BASF S.A. | Fluxapyroxad, pyraclostrobin |
| Priori | Fungicide | Syngenta | Azoxystrobin |
| Priori Xtra | Fungicide | Syngenta | Azoxystrobin, cyproconazole |
| Tilt | Fungicide | Syngenta | Propiconazole |
| **HERBICIDES** | | | |
| Ally | Herbicide | Du Pont | Metsulfuron-methyl |
| Aminol 806 | Herbicide | Adama Agricultural Solutions Ltd. | 2,4-D dimethylamine, acid equivalent of 2,4-D |

| | | | |
|---|---|---|---|
| Atectra | Herbicide | BASF | Dicamba |
| Heat | Herbicide | BASF | Saflufenacil |
| Topik 240 EC | Herbicide | Syngenta | Clodinafop-propargyl |
| **INSECTICIDES** | | | |
| Cruiser 350 FS | Insecticide | Syngenta | Thiamethoxam |
| Engeo Pleno | Insecticide | Syngenta | Thiamethoxam, lambda-cyhalothrin |
| Fastac Duo | Insecticide | BASF | Acetamiprid, alpha-cypermethrin |
| Imunit | Insecticide | BASF | Alpha-cypermethrin, teflubenzuron |
| Nomolt 150 | Insecticide | BASF | Teflubenzuron |
| Pirate | Insecticide, acaricide | BASF | Chlorfenapyr |
| Standak Top | Insecticide, fungicide | | Pyraclostrobin, thiophanate-methyl, fipronil |
| **ADJUVANTS** | | | |
| Aureo | Adjuvant | Bayer S.A. | Soybean oil methyl ester |
| Assist | Adjuvant | BASF S.A. | Mineral oil |
| Nimbus | Adjuvant | Syngenta Proteção de Culturas LTDA | Mineral oil |
| **GROWTH REGULATOR** | | | |
| Moddus | Growth regulator | Syngenta Proteção de Culturas LTDA | Trinexapac-ethyl |
