## Supplemental Table 2 for "Population structure determined by comparative amplicon gene sequencing in biofilm and planktonic bacterial communities from pesticide-contaminated water"

| Functional characteristics | r2 | P |
|---|---|---|
| Communities M1 and M2 | | |
| Motility | 0.6196 | <0.002 |
| Community B2 | | |
| Cellular processes | | |
| Growth and death | 0.8670 | <0.001 |
| Transportation in catabolism | 0.8977 | <0.001 |
| Environmental processes | | |
| Membrane transport | 0.9827 | <0.001 |
| Interaction of signaling molecules | 0.9422 | <0.001 |
| Signal transduction | 0.9838 | <0.001 |
| Genetic information processes | | |
| Translation | 0.9880 | <0.001 |
| Transcription | 0.9863 | <0.001 |
| Repair and replication | 0.9887 | <0.001 |
| Folding, sorting and degradation | 0.9888 | <0.001 |
| Metabolic processes | | |
| Enzymes | 0.9838 | <0.001 |
| Amino acids | 0.9872 | <0.001 |
| Secondary metabolites | 0.9883 | <0.001 |
| Carbohydrates | 0.9874 | <0.001 |

| | | |
|---|---|---|
| Energy | 0.9864 | <0.001 |
| Glycans | 0.9825 | <0.001 |
| Lipids | 0.9850 | <0.001 |
| Cofactors and vitamins | 0.9872 | <0.001 |
| Other amino acids | 0.9883 | <0.001 |
| Terpenoids, polyketides | 0.9845 | <0.001 |
| Nucleotides | 0.9865 | <0.001 |
| Xenobiotic biodegradation | 0.9402 | <0.001 |
