## Supplementary material for "Population structure determined by comparative amplicon gene sequencing in biofilm and planktonic bacterial communities from pesticide-contaminated water": Cover Letter Plos One

Dear Dr. Ana Lopes,

Academic Editor of PLOS ONE,

Enclosed for your consideration is a Research Article entitled “Biotechnological potential determined by comparative amplicom gene sequencing in biofilm bacterial communities from pesticide contaminated water”, at PLOS ONE.

The reasons why we believe it deserves to be published stem from the following features:

This study analyzed the bacterial population structure in planktonic and biofilm lifestyles in tanks where pesticide contaminated water was stored for six months, as well as possible adaptive strategies using the data originated from next-generation sequencing of the 16S rRNA gene. We found a higher variation in population structure and diversity in the biofilm than planktonic samples, indicating a possible combined genetic drift process followed by the selection of individuals surviving to stressful water conditions, as heating and contact with agrochemicals during a six months period. Possibly, the biofilm formation in water tanks contaminated with pesticides enabled the conditions for survival of the bacteria genera which has been harboring the essential processes for adaptation to these environments, according to description obtained from gene bank database. This study suggests the potential for adaptation to contaminated environments, degradation and bioremediation of pesticides by bacteria genera identified by sequencing of DNA from biofilm, which herbicide biodegradation kinetics can be exploited by culturable collection obtained from the communities studied in this work.

The authors declare no competing financial interests.

This article is not being considered for publication in another journal and it is the first submission at PLOS ONE.

Thank you for your time and consideration. I look forward hearing from you.

Sincerely yours,

Prof. Dr. Marcos Pileggi
