## Supplementary material for "Population structure determined by comparative amplicon gene sequencing in biofilm and planktonic bacterial communities from pesticide-contaminated water": English Review Certificate

AJE AMERICAN JOURNAL EXPERTS

### EDITORIAL CERTIFICATE

This document certifies that the manuscript listed below was edited for proper English language, grammar, punctuation, spelling, and overall style by one or more of the highly qualified native English speaking editors at American Journal Experts.

#### Manuscript title:

Populational structure determined by comparative amplicon gene sequencing in biofilm and planktonic bacterial communities from pesticide contaminated water

#### Authors:

Jhenifer Yonara Lima, Cassiano Moreira, Gessica Costa, Paloma Nathane Nunes Freitas, Luiz Ricardo Olchanheski, Sônia Alvim Veiga Pileggi, Rafael Mazer Etto, Christopher Staley, Michael Jay Sadowski, Marcos Pileggi

#### Date Issued:

July 5, 2019

#### Certificate Verification Key:

5085-76BC-AB9B-9531-ECBP

This certificate may be verified at www.aje.com/certificate. This document certifies that the manuscript listed above was edited for proper English language, grammar, punctuation, spelling, and overall style by one or more of the highly qualified native English speaking editors at American Journal Experts. Neither the research content nor the authors' intentions were altered in any way during the editing process. Documents receiving this certification should be English-ready for publication; however, the author has the ability to accept or reject our suggestions and changes. To verify the final AJE edited version, please visit our verification page. If you have any questions or concerns about this edited document, please contact American Journal Experts at.

*American Journal Experts provides a range of editing, translation and manuscript services for researchers and publishers around the world. Our top-quality PhD editors are all native English speakers from America's top universities. Our editors come from nearly every research field and possess the highest qualifications to edit research manuscripts written by non-native English speakers. For more information about our company, services and partner discounts, please visit www.aje.com.*
